# Differentiation of human intestinal organoids with endogenous vascular endothelial cells

**DOI:** 10.1101/2020.03.15.991950

**Authors:** Emily M. Holloway, Joshua H. Wu, Michael Czerwinkski, Caden W. Sweet, Angeline Wu, Yu-Hwai Tsai, Sha Huang, Amy E. Stoddard, Meghan M. Capeling, Ian Glass, Jason R. Spence

## Abstract

Human pluripotent stem cell (hPSC)-derived intestinal organoids (HIOs) generated using directed differentiation lack some cellular populations found in the native organ, including vasculature. Using single cell RNA sequencing (scRNAseq), we have identified a transient population of endothelial cells (ECs) present early in HIO differentiation that are lost over time in culture. Here, we have developed a method to enhance co-differentiation and maintenance of ECs within HIOs (vHIOs). Given that ECs are known to possess organ specific gene expression, morphology and function, we used bulk RNAseq and scRNAseq to interrogate the developing human intestine, lung, and kidney in order to identify organ-enriched EC-gene signatures in these organ systems. By comparing organ-specific gene signatures along with markers validated by fluorescent *in situ* hybridization to HIO ECs, we find that HIO ECs grown *in vitro* share the highest similarity with native intestinal ECs relative to kidney and lung. Together, these data show that HIOs can co-differentiate a native EC population that are properly patterned with an intestine-specific EC transcriptional signature *in vitro.*

## INTRODUCTION

The development of human pluripotent stem cell (hPSC) derived small intestinal organoids (HIOs) (Spence et al., 2010) or primary organoids derived from human donor tissues (Sato et al., 2011), has led to an improved understanding of human intestinal development and physiology. In order to generate hPSC-derived HIOs, recombinant proteins and/or small molecules are exogenously supplied to cultures in order to mimic the *in vivo* signaling environment present during development, through a process called directed differentiation (McCracken et al., 2011). Current methods to differentiate HIOs generate both endodermal cells that will give rise to the HIO epithelium as well as a mesoderm population that can give rise to cells of the lamina propria, such as fibroblasts and smooth muscle cells (Finkbeiner et al., 2015a, 2015b; McCracken et al., 2011; Spence et al., 2010; Watson et al., 2014; Wells and Spence, 2014). However, HIOs do not fully recapitulate the complexity of the native human intestine, lacking cellular components including immune cells, enteric neurons (Workman et al., 2016), vasculature and the microbiome (Hill et al., 2017a; Leslie et al., 2014).

Co-culture and transplantation approaches have been developed to increase the complexity of HIOs (Holloway et al., 2019). For example, hPSC-derived enteric neural lineages have been added *in vitro* and further matured *in vivo* to establish a functional enteric nervous-like system within HIOs (Schlieve et al., 2017; Workman et al., 2016). Similarly, microinjection of *E. coli* into the HIO lumen has permitted the study of epithelial response to early gut colonization (Hill et al., 2017a, 2017b). However, vascularization of HIOs has been restricted to *in vivo* models, whereby HIOs are transplanted into highly vascularized regions of immunocompromised mice (Cortez et al., 2018; Watson et al., 2014). In these environments, HIOs undergo extensive vascularization by the murine host tissue, and increase in complexity to resemble mature intestinal tissue (Finkbeiner et al., 2015b; Watson et al., 2014). However, co-culture approaches or co-differentiating HIOs with a native vasculature prior to *in vivo* engraftment has not yet been achieved.

Here, we performed single cell RNA sequencing (scRNAseq) at various timepoints across HIO differentiation *in vitro* and observed a transient population of endothelial-like cells (ECs) present within HIOs early during differentiation; however, these cells are not maintained during prolonged culture under standard growth conditions. This suggested that early during HIO differentiation, cells within the culture are capable of giving rise to EC-like cells. Based on these observations, we hypothesized that a modified directed differentiation approach would allow the induction and maintenance of a more robust EC population within HIOs (termed vHIO). Our findings demonstrate that modified culture conditions allow a ∼13-fold increase in the induction of EC-like cells within HIOs without impacting the other HIO cell populations present (i.e. epithelium, mesenchyme), and support the survival of this population of ECs within HIOs in culture for months.

Since organ-specific morphology in vascular beds has long been appreciated (Aird, 2007), and organ-specific transcriptional signatures and functions have been described in mouse (Ding et al., 2011; Kalucka et al., 2020; Lee et al., 2014; Nolan et al., 2013) and human tissues (Chi et al., 2003; Marcu et al., 2018), we further sought to determine if HIO ECs were properly patterned by interrogating human fetal intestine, lung, and kidney tissue to identify and validate organ enriched EC gene signatures and individual genes. These data were then used to assess the extent to which HIO ECs resembled human intestinal ECs. After two months of culture, we observed that relative to human fetal intestine, lung and kidney ECs, HIO ECs were the most transcriptionally similar to the fetal intestine ECs, suggesting that HIOs possess intrinsic properties sufficient to induce ECs to undergo proper organ-specific patterning.

Taken together, this work shows that a native EC population can be co-differentiated within HIO cultures, and that these EC-like cells can be maintained under defined culture conditions (vHIOs) for prolonged culture periods. We further present human intestinal, lung and kidney EC data sets that can be used as a reference for comparison against *in vitro* derived organoids and specific cell types found within *in vitro* organoids. Using transcriptional profiles and a panel of several validated markers, we conclude that HIO ECs undergo organ-specific patterning *in vitro*, and most closely resemble an intestinal EC population.

## RESULTS

### hPSC-derived human intestinal organoids (HIOs) develop a transient population of endothelial cells (ECs)

Human intestinal organoids (HIOs) have been almost exclusively characterized following growth for several weeks in culture (Capeling et al., 2019; Finkbeiner et al., 2015b; Spence et al., 2010; Tsai et al., 2017). Through these analyses, various intestinal epithelial and mesenchymal populations have been identified. Initial observations suggested that HIOs lacked an EC population (Spence et al., 2010); however, a thorough investigation of cellular heterogeneity within HIOs across time has not been carried out.

In order to gain insights into cellular heterogeneity during HIO growth and differentiation, we performed a single cell RNA sequencing (scRNAseq) time course analysis. Analysis was carried out on HIO samples following hindgut patterning and specification into a CDX2+ intestinal lineage, the time when monolayer cultures give rise to 3-dimensional (3D) spheroids (herein referred to as ‘day 0’), and several time points after embedding spheroids in Matrigel for growth and expansion into HIOs (days 3, 7, and 14) (Figure 1A). A total of 13,289 cells from these four time-points (days 0, 3, 7, 14) were included in the computational analysis and the data were visualized using UMAP dimensional reduction (Becht et al., 2019; Wolf et al., 2018) (Figure 1B). Cell classifications were carried out using canonical markers for lineages (i.e. epithelial, mesenchymal, endothelial) (Figure 1B, S1). We identified expected epithelial and mesenchymal cell lineages within HIOs; however, we also identified cell clusters possessing endothelial (cluster 14) and neuronal (clusters 11,15) gene signatures (Figure S1). The endothelial cluster was defined by a gene set expected to be present in ECs, including *CDH5, KDR, FLT1*, and *ESAM* (Figure 1C, S1). The time course revealed that the EC-like population in cluster 14 was transient, with the highest proportion of EC-like cells on day 3 of culture (1.35% of all day 3 cells), after which EC prevalence progressively decline to 0.54% on day 7 and 0.42% on day 14 (Figure 1D). Wholemount staining of day 3 HIOs with the EC lineage markers CD144 and CD31 confirmed the existence of an EC-like population early during HIO growth (Figure 1E). Together, these data suggest that HIO differentiation cultures are capable of spontaneously giving rise to EC-like cells that diminish over time.

**Figure 1.**
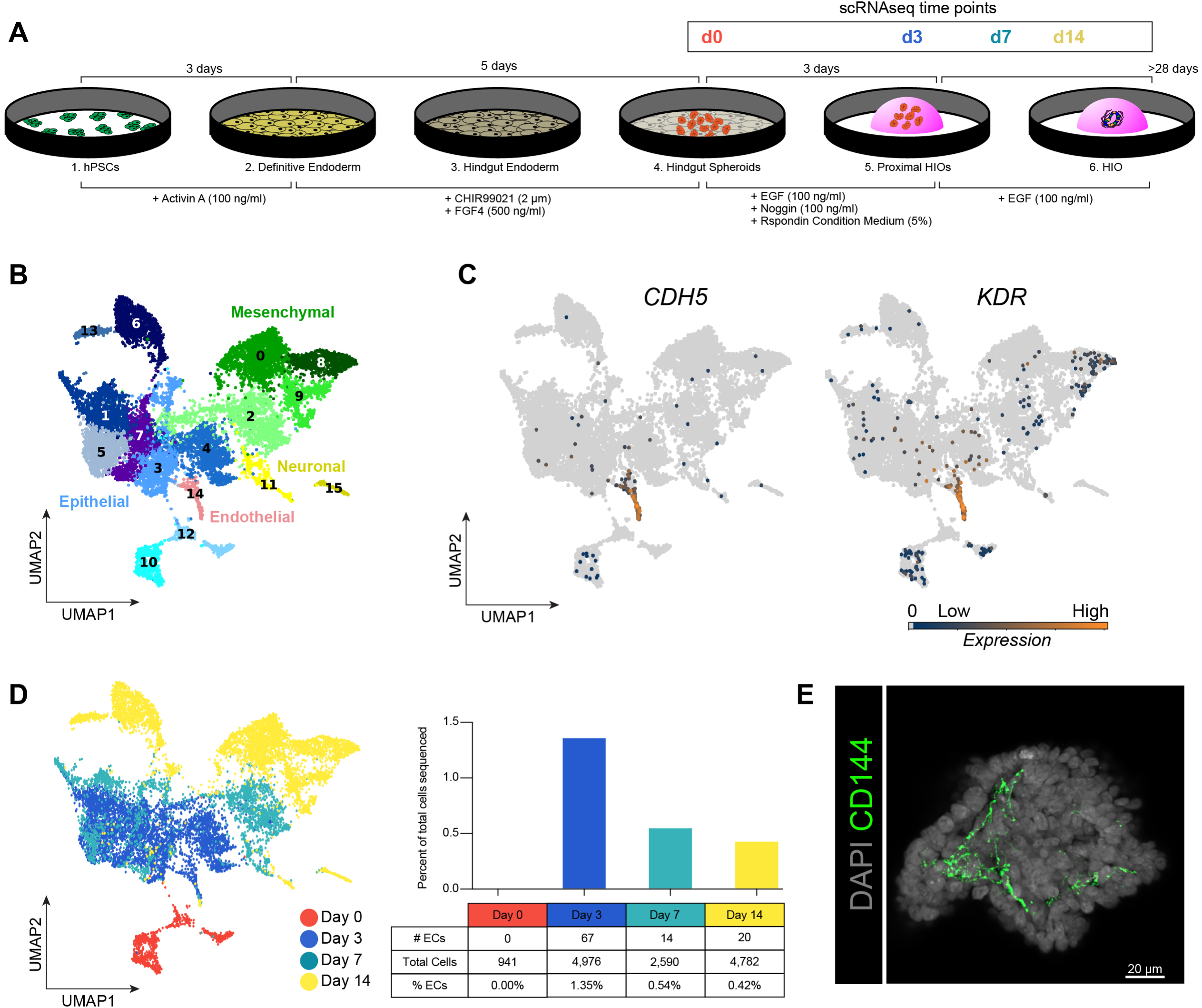
Identification of an endothelial cell-like population in HIOs. **(A)** Overview of HIO differentiation protocol highlighting time points when HIOs were collected for single-cell RNA sequencing (scRNAseq). **(B)** UMAP plot of 13,289 cells from all time points profiled with scRNAseq predicted 15 cell clusters. Cluster identities were assigned based on expression of canonical lineage markers for epithelium, mesenchyme, endothelium, and neurons (see also Figure S1). **(C)** Feature plots for endothelial cell (EC) markers *CDH5* and KDR showing enrichment in Cluster 14. **(D)** UMAP plot illustrating the distribution of cells colored by sample (time point) and demonstrating the proportion of Cluster 14 (putative ECs) relative to total cells sequenced per timepoint (bar chart). **(E)** Wholemount immunofluorescent staining of d4 HIOs with the EC marker CD144 (green) and DAPI (grey). Scalebar represents 20 µm.

### Developing a method to induce and maintain ECs within HIOs

The progressive reduction of EC-like cells over time suggested that standard HIO culture conditions are not optimal for supporting robust long-term EC survival. Therefore, we sought to define new culture conditions that would increase the survival of the co-differentiated EC population within HIOs. We hypothesized that addition of growth factors important for vascular induction and maintenance would improve EC differentiation and maintenance within HIOs, including VEGF, bFGF, and BMP4 (Orlova et al., 2014; Patsch et al., 2015; Sriram et al., 2015; Wimmer et al., 2019). On day 2 of HIO culture, VEGF was added into the HIO media (EGF/NOG/RSPO – ‘ENR’) in order to promote survival of any endogenously arising ECs (Figure 2A). Although BMP has been used in multiple culture settings to induce ECs from hPSCs, NOG was included for the first 3 days of HIO growth because modulation of BMP signaling is also responsible for proximal-distal patterning in HIOs (Múnera et al., 2017). Antagonism via NOG is required for patterning HIOs into proximal duodenum-like tissue, while adding BMP ligands to the media in early cultures instructs a colonic fate in HIOs (Múnera et al., 2017). After 3 days of HIO patterning into an proximal/duodenal identity, HIOs were maintained without RSPO or NOG from day 3 onwards, as EGF is sufficient to support HIO growth following proximal-distal patterning (Múnera et al., 2017). Subsequently, in order to induce additional ECs, HIOs were treated for three days with VEGF, FGF2 (bFGF) and BMP4 (herein referred to as the vHIO protocol; Figure 2A), which have been shown to enrich ECs in hPSC-derived mesenchyme (Orlova et al., 2014; Patsch et al., 2015; Sriram et al., 2015; Wimmer et al., 2019). HIOs generated with the vHIO protocol were maintained in media supplemented with EGF and VEGF for up to two months prior to analysis, while controls were maintained in media containing EGF only (Figure 2A). To assess the proportion of EC-like cells in HIOs following induction, flow cytometric analysis of CD31+/CD144+ cells was used to compare control HIOs and vHIOs. After several weeks in culture, ECs constituted an average of 0.20% in control HIOs, whereas vHIOs possessed 2.62% CD31+/CD144+ a ∼13-fold increase compared to control culture conditions (Figure 2B). Identification of CD31+/CD144+ cells within vHIOs using immunofluorescence confirmed EC enrichment as double-positive cells were abundant and were observed within HIO mesenchyme (Figure 2C, S2C). ECs persisted for two months in culture, the longest time point examined (Figure 2D,E, S2E).

**Figure 2.**
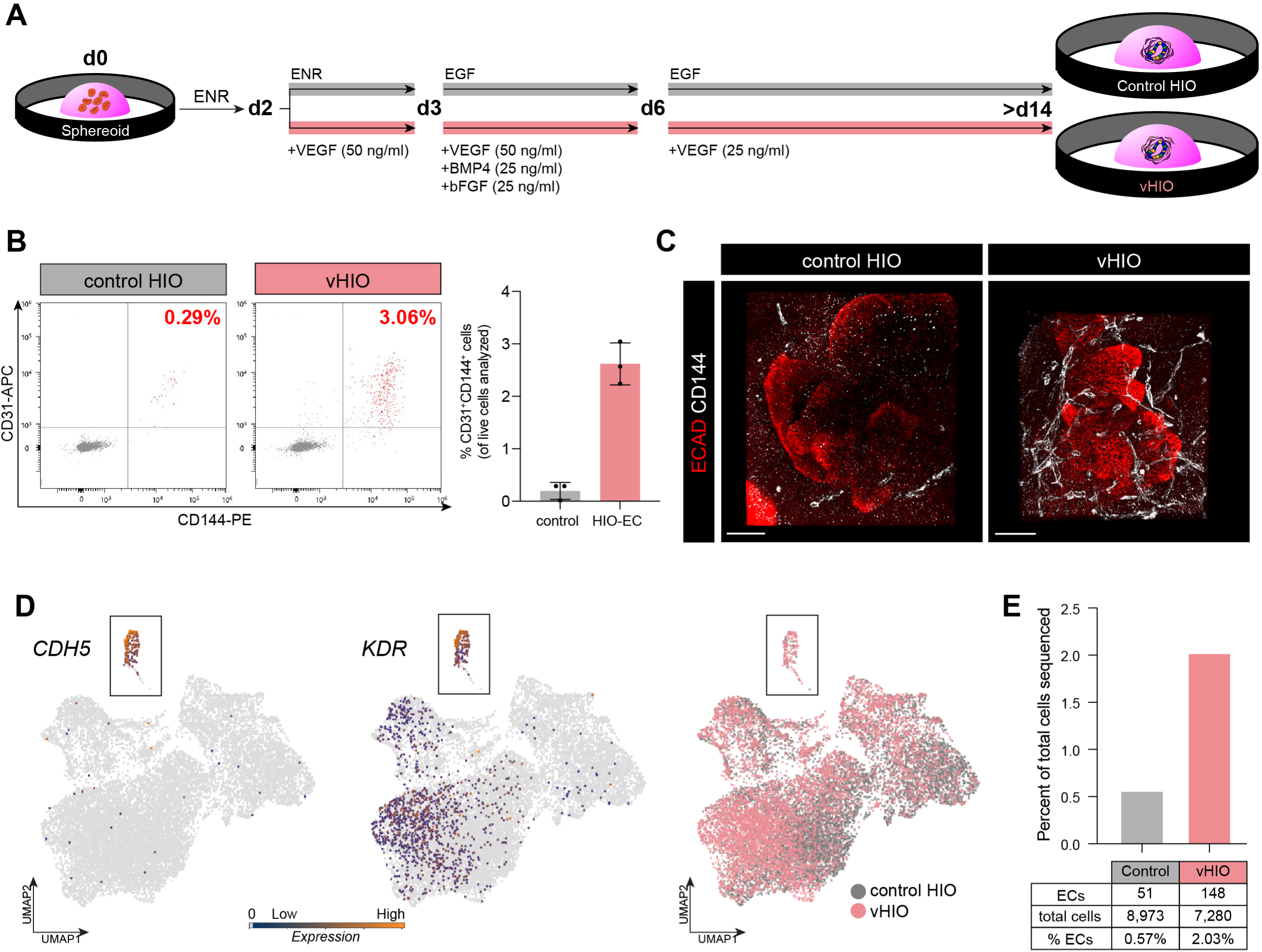
A method to increase differentiation and survival of ECs in HIOs. **(A)** Schematic of control and vHIO differentiation paradigms. **(B)** Representative flow cytometric plots of control HIOs and vHIOs for EC markers CD31 and CD144. Flow cytometric analysis of independent batch-matched differentiations of control HIO and vHIOs for presence of CD31+/CD144+ cells after 2-3 weeks in culture (n=3 differentiations in 2 different cell lines), showing an average ∼13-fold increase of EC-like cells in the vHIOs. **(C)** Maximum intensity projection of a wholemount confocal z-series staining for the EC marker CD144 (white) along with ECAD (red) in control HIOs and vHIOs. Scale bar represents 100 µm. (D) UMAP plots demonstrating scRNAseq data from 59d control HIO (grey) and vHIO (pink), and showing feature plots of the EC markers *CDH5* and *KDR* (total of 16,253 cells). Boxed region around the putative EC Cluster 5 (See also Figure S2). (E) Relative proportion of EC-like cells in control HIOs and vHIO conditions compared to the total number of cells sequenced after 59d in culture.

To further support our flow cytometric and immunohistochemical data showing EC-enrichment, and to assess the effect of the vHIO protocol on overall HIO development and cellular makeup, scRNAseq was performed on control and vHIO tissue after two months in culture. A total of 16,253 cells (8,973 control and 7,280 vHIO) were analyzed and the data were visualized using UMAP dimensional reduction (Figure 2D-E). Epithelial lineages (clusters 2 and 3), mesenchymal lineages (clusters 0,1, and 4), and endothelial lineages (cluster 5) were classified based on expression of canonical makers (Figures 2D and S2D-E). Quantification of the contribution of control and vHIO cells to each cluster was performed (Figure S2D). ECs were enriched in the vHIO condition compared to control, comprising 2.03% of all cells sequenced in the vHIO condition, a 3.5-fold increase over control HIOs (Figure 2E). More generally, this scRNAseq analysis demonstrated that all of the cell populations visualized by UMAP have contributions from both HIOs and vHIOs, showing that the vHIO differentiation paradigm does not induce cell population gain or loss (Figure S2D), and reinforces data showing that this method enriches EC-like cells (Figure S2D).

### Defining a human intestinal EC transcriptional signature

Organ-specific properties including morphology, transcriptional signatures, and function have been described in mouse and human organs (Daniel et al., 2018; Ding et al., 2010, 2011; Lee et al., 2014; Marcu et al., 2018; Nolan et al., 2013); however, we sought to better understand intestine-specific EC properties in order to compare *in vivo* ECs with induced ECs found within HIOs. To identify intestine-specific EC transcriptional signatures, we performed bulk RNA sequencing (RNAseq) on FACS-isolated EC (CD31^+^CD144^+^) and non-EC (CD31^-^CD144^-^) populations from human fetal intestine, lung, and kidney spanning 13-20 weeks of development (Figure 3A). Bulk RNAseq data demonstrated that canonical EC markers (*CD31, CD144, KDR*) were enriched in the CD31+/CD144+ isolated populations (Figure 3B). Additionally, the EC samples had low-to-no expression of mesenchyme genes (i.e. *PDGFRa*), hematopoietic genes (i.e. *CD45*), and epithelial genes (i.e *EPCAM)*, which were expressed in the double negative (CD31^-^CD144^-^) population as expected (Figure 3B). Unsupervised hierarchical clustering of all samples revealed that primary ECs from different human organs (intestine, lung, kidney) formed their own clade, and the CD31-/CD144-‘non-ECs’ formed another clade. Notably, biological replicates of the same organ were closest in similarity compared to EC samples isolated from the other organs, to non-ECs, and to human umbilical vein endothelial cells (HUVECs), suggesting that there are organ-specific transcriptional profiles among human fetal ECs (Figure 3C). Principal component analysis of the primary ECs produced organ-specific clustering, further supporting the existence of organ-specific transcriptional differences across human fetal intestine, lung, and kidney ECs (Figure 3D). K-means (k=20) gene clustering was used to identify genes enriched within the ECs of a single organ. At this resolution, pan-EC enriched gene clusters (0, 1, 2, 3, 4) as well as organ-specific EC-enriched gene clusters (6, 8, 17) were identified (Figure 3E). Over 100 organ-specific EC-enriched genes were computationally identified for all 3 organs. These gene lists are herein referred to as the lung EC signature (lECs), intestine EC signature (iECs), and kidney EC signature (kECs) (Supplemental Table 1).

**Figure 3.**
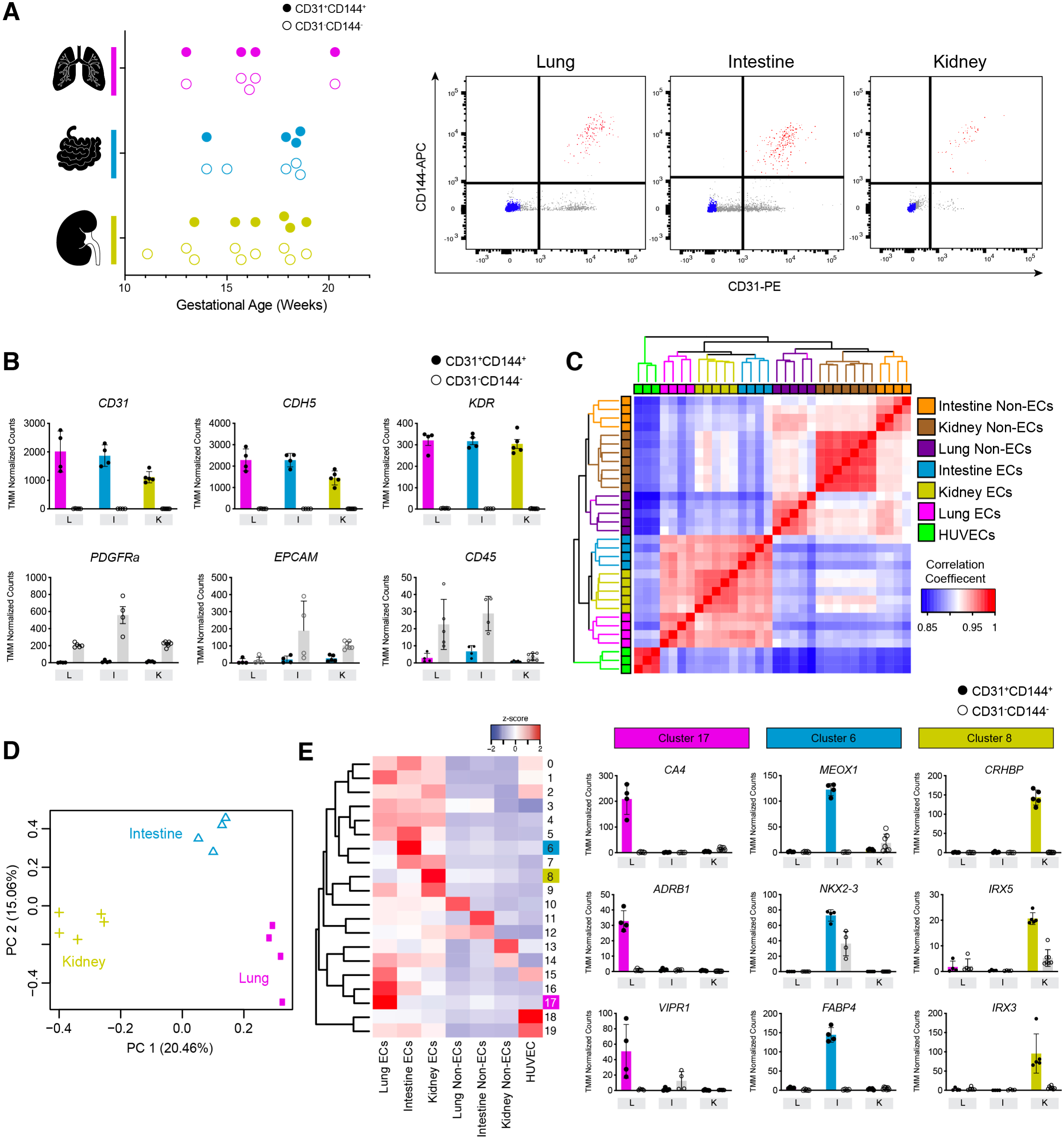
Defining organ-specific EC genes and gene signatures during human development. **(A)** Left: Summary of all human lung, intestine, and kidney samples profiled by bulk RNAseq. Closed circles represent samples collected from sorted EC populations, and open circles represent sorted non-EC populations. Samples are colored by organ system: Pink – lung; Blue – intestine; Yellow – kidney. Right: representative flow cytometry plots from each organ. ECs (CD31+CD144+, red) and non-ECs (CD31-CD144-, blue) populations were isolated using FACS. **(b)** Bulk RNAseq data showing TMM normalized counts for EC genes (*CD31, CDH5, KDR*) highly enriched in the CD31+CD144+ samples, and non-EC genes associated with the mesenchyme (*PDGFRa*), epithelial (*EPCAM*), and immune (*CD45)* transcripts enriched in CD31-CD144-samples. **(C)** Unsupervised hierarchical clustering of all samples profiled in this analysis, along with HUVECs. Each row corresponds to a biological replicate (except HUVEC samples, which represent n=3 technical replicates). **(D)** Principal component analysis of primary EC populations. **(E)** Left: K-means gene clustering (k=20, labeled 0-19) identifies organ-specific EC-enriched genes. Each row represents a cohort of genes enriched in one or more samples. Expression of the cohort is shown in each sample as the average normalized gene expression of all genes in that group. Intestinal EC-enriched genes (Cluster 6 – blue box), Kidney EC-enriched genes (Cluster 8 – yellow box), Lung EC-enriched genes (Cluster 17 – pink box). Right: TMM normalized expression of representative organ-specific EC enriched gene candidates in human lung (pink: *CA4, ADRB1, VIRP1)*, intestine (blue: *MEOX1, NKX2.3, FABP4*), and kidney (yellow: *CRHBP, IRX5, IRX3*) are shown across all primary samples. Closed circles represent samples collected from sorted EC populations, and open circles represent sorted non-EC populations.

Validation of the organ-specific EC genes and gene signatures identified by bulk RNAseq (Figure 3) was carried out using both scRNAseq and *fluorescent in situ hybridization* (FISH) (Figure 4). We used scRNAseq to profile human fetal lung (Miller et al., 2020), intestine (Czerwinski et al., 2020), and kidney across 7 specimens spanning 13.5-19 weeks of gestation (Figure S4A). In total, our analyses included 62,046 cells across these three organs (Figure 3A and S3A). An EC cluster was identified for each organ using cell type scoring (see Methods), which leverages gene cohorts canonically associated with different cell classes (i.e. epithelium, mesenchyme, endothelium, immune, neuronal) (Miller et al., 2020) (Figure 4A, S4). Based on this analysis, EC clusters were computationally extracted and re-clustered; the EC clusters collectively contained 3,082 cells and are comprised of 1,361 lung ECs, 877 intestine ECs, and 884 kidney ECs (Figure 4B, Figure S3A). This analysis also revealed EC heterogeneity in the form of sub-clusters, including a lymphatic EC cluster apparent in the lung (cluster 5) and intestine (cluster 2) as defined by *PROX1* expression (De Val and Black, 2009; Wigle and Oliver, 1999; Wigle et al., 2002) (Figure S3B). Organ-specific EC genes identified in bulk RNAseq data were validated in scRNAseq data (Figure 4C, E, and G, Figure S3C-E). The single cell data showed that organ-specific EC genes are expressed by vascular ECs in a manner expected based on the bulk sequencing data. Notably, despite being enriched in ECs in an organ-specific manner, markers were also expressed by non-EC lineages among the organs profiled (Figure S4). Collectively, these data showed that a given gene is enriched in ECs of a single organ relative to the other organs, but that expression may not be exclusive to ECs (Figure S4). We also used multiplexed FISH to validate a cohort of organ-specific EC genes. Sample matched human fetal lung, intestine, and kidney were stained for the panel of organ-specific EC markers alongside the pan-EC marker *CDH5* (Figure 4D, F, H and S3). Three EC-specific organ-enriched genes were selected from lung (*CA4, ADRB1, VIRP1*), intestine (*MEOX1, NKX2.3*, FABP4) and kidney (*CRHBP, IRX3, IRX5*) for validation by multiplexed FISH (Figure 3E, Figure 4). This approach confirmed data obtained in bulk and single cell RNA sequencing. Through this validation effort, a panel of 9 organ-specific EC enriched markers across human fetal lung, intestine, and kidney can be used in combination with computational analysis to assess the patterning of ECs co-differentiated within HIOs.

**Figure 4.**
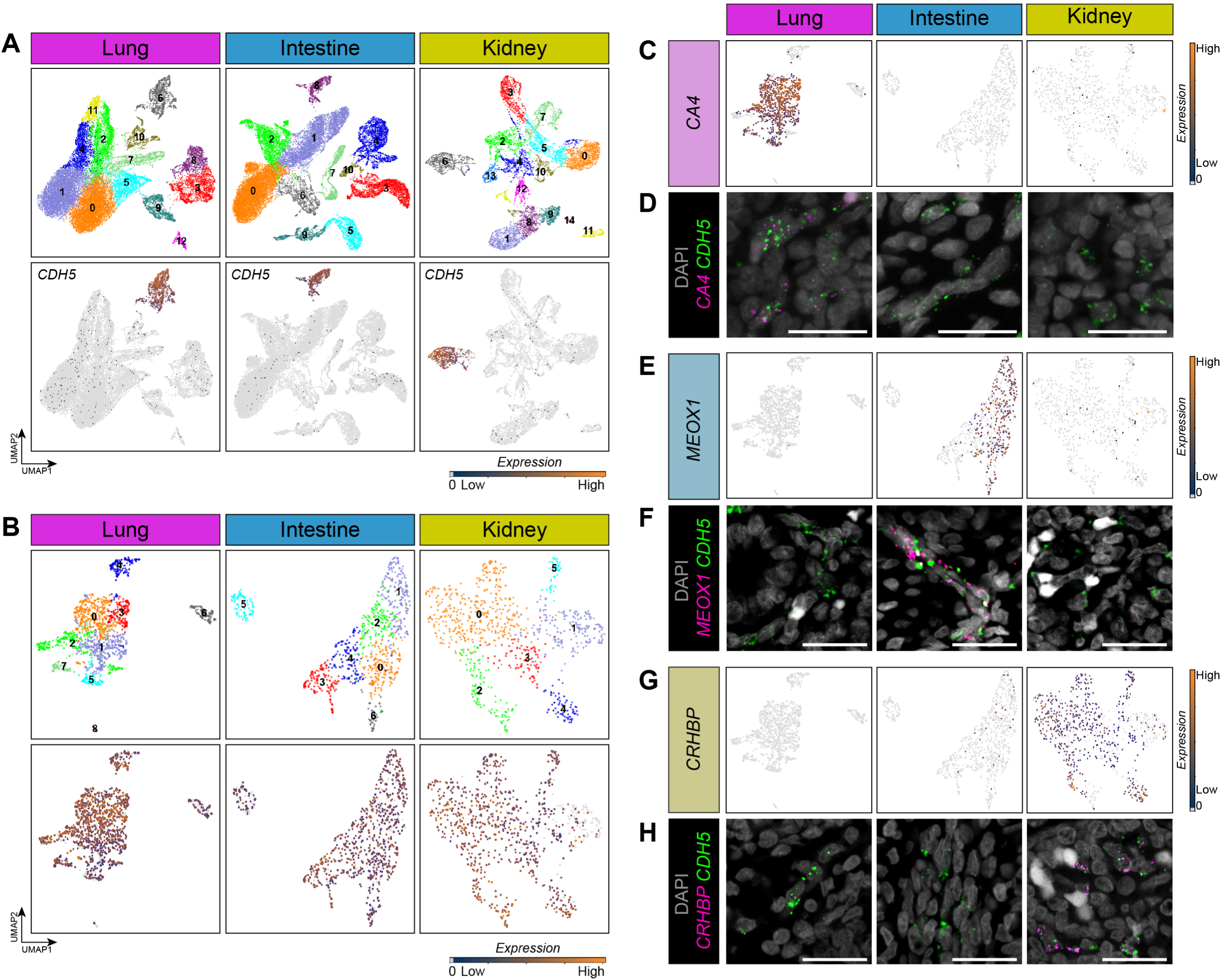
Validation of organ-specific EC-enriched genes. **(A)** Top row: UMAP plots for human lung (Pink; n= 3; 26,501 cells), intestine (Blue; n=6; 26,010 cells), and kidney (Yellow; n=2; 9,535 cells) showing predicted cell clusters. Bottom row: Feature plots for the EC-specific marker *CDH5*, highlighting the EC population within each organ. **(B)** Top row: EC clusters from each organ were computationally extracted, re-clustered, and visualized using UMAP. A total of 1,361 lung ECs, 877 intestinal ECs, and 844 kidney ECs were included in the analysis. Bottom row: Feature plots of the EC marker CDH5 among extracted clusters for each organ. **(C-D)** C: Feature plots of organ-specific EC genes (identified in Figure 3) showing expression in scRNA-seq data of the lung-specific EC candidate *CA4* in ECs of the lung, intestine and kidney. D: Multiplexed FISH for the pan-EC marker *CDH5* (green) and *CA4* (pink) in sample matched human fetal tissue (117d). **(E-F)** E: Feature plot of scRNA-seq data of the intestine-specific EC candidate *MEOX1* expression among primary ECs. F: Multiplexed FISH for the pan-EC marker *CDH5* (green) and *MEOX1* (pink) in sample matched human fetal tissue (117d). **(G-H)** G: Feature plots of scRNA-seq data of the kidney-specific EC candidate *CRHBP* expression among primary ECs. H: Multiplexed FISH for the pan-EC marker *CDH5* (green) and *CRHBP* (pink) in sample matched human fetal tissue (120d). Scalebars represent 25 µm.

### HIO ECs share the highest transcriptional similarity with fetal intestinal ECs

Given the unique organ-specific gene expression by ECs *in vivo* during human development, we wanted to assess the extent of organ-specific patterning of the ECs induced within HIOs. The EC cluster from scRNAseq (cluster 5, Figure S2D) of 59d control and HIO ECs was computationally extracted, reclustered and visualized using UMAP (Figure 5A). To determine how similar gene expression of HIO ECs was to native organ gene signatures, organ-specific EC-enriched gene signatures (lECs, iECs, and kECs) identified in bulk RNAseq (Figure 3) served as the foundation for organ-specific cell type scoring analysis. The HIO ECs were analyzed for expressed genes found in lECs, iECS, and kECs. As such, the expression of 137 lECs, 89 iECs, and 50 kECs genes was determined in HIO ECs, and the average expression for each organ-signature was determined for HIO ECs. Based on this analysis ECs co-differentiated within HIOs had the highest average expression of the intestinal EC signature relative to lung and kidney ECs (Figure 5B,C). FISH analysis of 29-day vHIOs confirmed expression of the intestine-specific marker, *NKX2.3* by HIO ECs (Figure 5D); and a lack of detectable expression of markers validated for the lung or kidney (Figure S5). Notably, the intestinal EC marker *MEOX1* was not detectable by FISH in vHIOs, suggesting that intestinal patterning may be incomplete in HIO derived ECs at this early time point (Figure S5).

**Figure 5.**
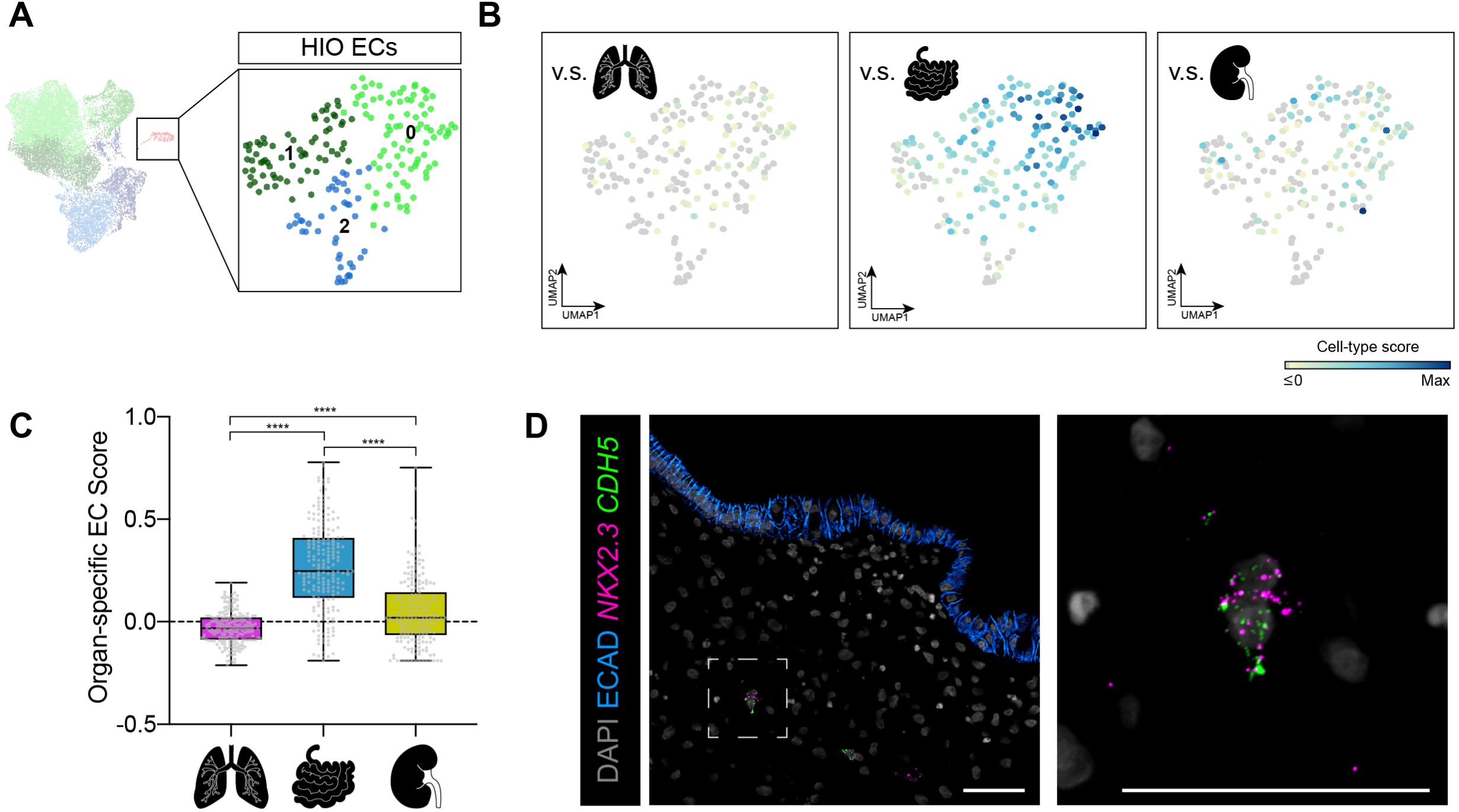
HIO ECs are transcriptionally similar to primary intestinal ECs. **(A)** The EC cluster from Control HIO and vHIO scRNA-seq data (as shown in Figure 2) was computationally extracted, re-clustered and visualized using UMAP. **(B)** Expression of genes identified in gene signatures from primary human fetal ECs (as identified in Figure 3) were interrogated in ECs from HIOs. A ‘gene signature score’ was calculated as the average expression of organ-specific EC genes (from primary tissue gene signatures) in the ECs of the HIO (see Methods). The score, corresponding to the average expression of the signature gene sets in each cell, is plotted. **(C)** Box-and-whiskers plot of individual data points shown in (B) of the organ-specific endothelial cell type scoring. Statistical significance was calculated using a one-way ANOVA (alpha = 0.05) followed by a Tukey post-hoc test. The intestine EC score was statistically different than both lung and kidney EC scores (p< 0.001). **(D)** FISH in an vHIO (29d) for the intestine-specific marker *NKX2.3* (pink) and the pan-EC marker *CDH5* (green), along with protein staining for ECAD (blue), and DAPI (grey). Scalebars represent 50 µm.

## DISCUSSION

While organoids are genetically tractable, complex *in vitro* human systems, they do not entirely recapitulate the full complement of cell types or complex physiology found in native tissues (Holloway et al., 2019). Several groups have been working to improve organoids, and approaches include implementing strategies to increase organoid complexity and maturation to more accurately mimic the native tissue (Fujii et al., 2018; Low et al., 2019; Mansour et al., 2018; Ouchi et al., 2019). In the case of cell types lacking in HIOs, such as ectoderm-derived enteric neurons, co-culture techniques using enteric neuron precursors (vagal neural crest cells) have been developed, facilitating the integration of a functional enteric nervous system that can stimulate motility following transplantation (Schlieve et al., 2017; Workman et al., 2016). While HIO vascularization has been achieved following *in vivo* transplantation by the host vasculature, developing a native human vasculature within HIOs has been elusive. Co-culturing endothelial cells (ECs) has proven successful in other *in vitro* systems, including hPSC-derived hepatic endoderm cultured with primary and hPSC-derived ECs leading to complex liver bud organoids with improved hepatocyte function (Camp et al., 2017; Takebe et al., 2013, 2017). A unique aspect of hPSC-derived HIOs is the co-differentiation of both intestinal epithelium and mesenchyme, which gives rise to diverse mesenchymal lineages including smooth muscle, myofibroblasts, and fibroblasts found in HIOs *in vitro* and following *in vivo* transplantation (Finkbeiner et al., 2015b; Spence et al., 2010; Watson et al., 2014). In this current work, we leveraged the plasticity of mesodermal progenitor cells early in HIO differentiation and demonstrated that a subset of these cells can be induced to differentiate into ECs.

Through a scRNAseq time course analysis of HIO development, we demonstrate that differentiation of HIO mesenchyme into vasculature takes place normally during early HIO differentiation, but that these cells are rare, and are mostly lost over time. Previous work has benchmarked d0 HIOs (intestinal spheroids) to embryonic day 8.5 (E8.5) mouse intestine (Spence et al., 2010). Vascularization of the gut is thought to begin a day later, around E9.5 (Hatch and Mukouyama, 2014), which follows a similar developmental progression as young HIOs, with ECs emerging within 72 hours of 3D culture. By targeting young HIOs with exogenous vascular induction and survival cues, we were able to increase the proportion and longevity of ECs within HIOs. While this is the first report demonstrating an endogenous EC population within HIOs, EC populations have been previously described in human kidney organoids (van den Berg et al., 2018; Combes et al., 2019; Czerniecki et al., 2018; Freedman et al., 2015; Low et al., 2019; Takasato et al., 2015). Vasculature is a mesoderm derivative (Ferguson et al., 2005; Risau and Flamme, 1995), although the exact origins of intestinal and renal ECs are not fully understood. The robust mesoderm patterning that occurs in both organoid systems likely produces precursors for the native EC populations observed; however, recent pseudotime analysis performed on human kidney organoids suggests that ECs might be derived from a subset of mesodermal nephron progenitor cells that co-express *KDR* (Low et al., 2019). Similar to what we have observed in HIOs, ECs within human kidney organoids can be expanded with targeted growth factor modulation (Czerniecki et al., 2018; Low et al., 2019) or exposure to flow using microfluidics (Homan et al., 2019). Incorporation of flow into the vHIOs is an exciting future direction for this platform, as it might expand and enhance the stability the vascular networks and increase the appeal of this system for drug discovery.

Several studies have demonstrated that vasculature patterning is organ-specific (Daniel et al., 2018; Ding et al., 2010; Feng et al., 2019; Kalucka et al., 2020; Lee et al., 2014; Marcu et al., 2018; Nolan et al., 2013); however, whether or not organoids are programmed with this patterning information *in vitro* was not known. Furthermore, profiling of organ-specific ECs has not included human intestinal ECs prior to this work. Our data define a human intestinal EC signature that includes almost 150 genes that are enriched in intestinal ECs relative to lung and kidney ECs. Recent work described an adult murine EC atlas spanning 11 organ systems at single cell resolution, and provides a foundational resource to compliment the work presented here, aimed at understanding human intestinal EC patterning (Kalucka et al., 2020). Several of the validated human organ-specific markers identified in our dataset demonstrated similar expression patterns in adult mouse atlas, suggesting that organ-specific signatures established during development are retained into adulthood, although future work should confirm these findings in both developing and adult human and murine tissue. Further, a formal interrogation into mouse-human EC differences will yield an important understanding of species-specific differences. A unique strength of the current approach is the simultaneous characterization of organ-matched EC and non-EC populations. This strategy facilitated the identification of organ-specific EC-enriched signatures while at the same time understanding where and when these markers may be expressed in non-EC cell types in other organ systems.

While the vHIO platform described here constitutes the first *in vitro* organoid model known to contain appropriately pattered endogenous vasculature, it remains unknown what cell types and signals are responsible for inducing organ-specific gene expression and patterning in ECs. However, the scRNAseq characterization of HIOs across developmental time provides an unprecedented insight into the diverse cell types present, and future work can leverage both scRNAseq and spatial transcriptomics to interrogate this patterning, and can leverage the modular nature of the HIO system to systematically test which cell type(s) and the molecular mechanisms that are responsible for inducing intestine-specific patterning in ECs. Better understanding these mechanisms will likely shed significant light on organ development. Reciprocal signaling has been shown to occur between ECs and the surrounding organ microenvironment during development (Kao et al., 2015; Lammert, 2001; Lammert et al., 2003; Lazarus et al., 2011; Matsumoto et al., 2001; Vila Ellis et al., 2020) and ECs can influence development through the supply of membrane-bound or secreted factors, termed “angiocrine factors” (Rafii et al., 2016). Identifying these angiocrine roles of ECs has been challenging to study using *in vivo* animal models, as the vascular is highly sensitive to modulations *in vivo* (Ferrara et al., 1996; Shalaby et al., 1995), and it is technically challenging to parse out unique angiocrine roles from metabolic requirements for the vasculature. Thus, the vHIO model comprises a novel *in vitro* system that can be used to study the dynamic EC-organ crosstalk during intestinal development. Future work will focus on both how ECs are instructed to adopt organ-specific patterning, and also how ECs, in a perfusion independent context, might contribute to HIO development and maturation *in vitro*.

Taken together, our data identified an unexpected, rare population of EC-like cells that arise early during HIO differentiation. By developing a method that incorporates EC-inductive and maintenance growth factors, this population can be further expanded and maintained for months *in vitro*. Through extensive transcriptional characterization and validation of organ-specific primary ECs across primary intestine, lung, and kidney during human development, we demonstrate the ECs co-differentiated within HIOs do undergo organ-specific patterning *in vitro*. vHIO improves both complexity and biological resemblance to the native developing intestine and comprises an amenable model to study EC-organ crosstalk during intestinal organogenesis.

### Financial Support

JRS is supported by the Intestinal Stem Cell Consortium (U01DK103141), a collaborative research project funded by the National Institute of Diabetes and Digestive and Kidney Diseases (NIDDK) and the National Institute of Allergy and Infectious Diseases (NIAID). JRS is also supported by the National Heart, Lung, and Blood Institute (NHLBI - R01HL119215), by the NIAID Novel Alternative Model Systems for Enteric Diseases (NAMSED) consortium (U19AI116482.) and by grant number CZF2019-002440 from the Chan Zuckerberg Initiative DAF, an advised fund of Silicon Valley Community Foundation. IAG and the University of Washington Laboratory of Developmental Biology was supported by NIH award number 5R24HD000836 from the Eunice Kennedy Shriver National Institute of Child Health and Human Development (NICHD). EMH was supported by the Training in Basic and Translational Digestive Sciences Training Grant (NIH-NIDDK 5T32DK094775), the Cellular Biotechnology Training Program Training Grant (NIH-NIGMS 2T32GM008353), and the Ruth L. Kirschstein Predoctoral Individual National Research Service Award (NIH-NHLBI F31HL146162). MC was supported by the Training Program in Organogenesis (NIH-NICHD T32 HD007505). MMC was supported by Cellular Biotechnology Training Program Training Grant (NIH-NIGMS 2T32GM008353) and the NSF-GRFP (DGE 1256260) Additional support was provided by the University of Michigan Center for Gastrointestinal Research (UMCGR) (NIDDK 5P30DK034933).

## Acknowledgements

We thank Judy Opp and the University of Michigan Advanced Genomics Core for their expertise operating the 10X Chromium single cell capture platform and sequencing expertise. We would also like to thank the University of Michigan Microscopy and Flow Cytometry cores for providing access to confocal microscopes and image analysis software and cytometers, respectively. We would also like to thank the University of Washington Laboratory of Developmental Biology staff.

## Author contributions

EMH and JRS conceived the study. JRS supervised the research. EMH developed tissue dissociation methods and isolated primary ECs for bulk RNA sequencing. AW, YHT, EMH, MC developed tissue dissociation methods and generated single cell RNA sequencing data. JW and MC performed computational analysis. MC, JW, EMH and JRS interpreted computational results. EMH, CS, and AES performed IF and FISH experiments and imaging. EMH, CWS, AES, and AW, and YHT processed primary tissue for IF/FISH. EMH, SH, and MMC generated human intestinal organoids. IG provided critical material resources for this work. EMH assembled figures. EMH and JRS wrote the manuscript. EMH, JW, AW, and MC contributed methods. All authors edited, read and approved the manuscript.

## METHODS

Lead contact and materials availability: Contact Jason R. Spence at for requests for materials.

### Human pluripotent stem cells

The University of Michigan Human Pluripotent Stem Cell Research Oversight Committee approved all experiments using human embryonic (ESC) and induced pluripotent stem cells (iPSC). HIOs were generated from 3 independent lines for these studies. Control organoids were generated from hESC line H9 (NIH registry no 0062) and iPSC lines WTC11 (Kreitzer et al., 2013) and 72.3 (McCracken et al., 2014).

### Derivation of human intestinal organoids from hPSCs

Differentiation of hPSCs into human intestinal organoids (HIOs) was carried out as previously described (Tsai et al., 2017). Briefly, hPSCs were patterned into definitive endoderm (DE) by supplementing Roswell Park Memorial Institute 1640 (RPMI-1640) with Activin A (100 ng/ml) for 3 days and increasing HyClone FBS concentration (0%, 0.2%, 2%) each day of DE induction. Hindgut patterning was carried out through addition of FGF4 (500 ng/ml) and CHIR99021 (2 µm) to RMPI-1640 containing 2% HyClone FBS. Media was changed daily, and spheroids were collected after 5 days of hindgut patterning. Spheroids were embedded in Matrigel (8 mg/ml, Corning, 354234) and incubated in ENR media (Minigut basal media, supplemented with 100 ng/ml EGF, 100 ng/ml Noggin, and 5% R-Spondin 2 conditioned media) for 3 days to pattern duodenal identity. After 3 days, ENR media was replaced with Minigut media containing only EGF (100 ng/ml). HIOs were analyzed between 14-28 days of culture. Minigut media is composed of the following components: Advanced DMEM:12 (Life Technologies, 12634), 1x B27 supplement (Life Technologies, 17504044), L-2 mM Glutamine (Life Technologies, 25030), 15 mM HEPES (Life Technologies, 15630080). All medias used in differentiation contain 1x PenStrep (Life Technologies, 15140).

To generate vHIOs, the same protocol as above was followed with the following modifications; VEGF (50 ngl/ml) was added to ENR media after 2 days of spheroid culture. The following day, ENR was replaced by Minigut containing EGF (100 ng/ml), VEGF (50 ng/ml), bFGF (25 ng/ml), BMP4 (25 ng/ml) for 3 days. After 3 days, vHIOs were grown in Minigut media supplemented with EGF (100 ng/ml) and VEGF (25 ng/ml) for the duration of culture. vHIO and matched control HIOs were generated from H9 and WTC11 hPSC lines.

### Primary tissue collection

Use of human tissue was reviewed and approved by The University of Michigan Institutional Review Board (IRB). De-identified human fetal lung, intestine, and kidney tissue was obtained from the University of Washington Laboratory of Developmental Biology. Tissue was shipped overnight in Belzer-UW Cold Storage Solution (ThermoFisher, NC0952695) with cold packs, as previously published (Menon et al., 2018; Miller et al., 2020). A list of tissue specimens can be found in the key resources table.

### Tissue fixation, paraffin processing and storage

Sample-matched human fetal lung, intestine, and kidney tissue samples were collected and processed into ∼1-2 cm fragments. Tissues were fixed for 24 hours at room temperature in 10% Neutral Buffered Formalin (NBF), and washed with UltraPure Distilled Water (Invitrogen, 10977-015) for 3 changes for a total of 2 hours. Tissue was dehydrated in a methanol series (25%, 50%, 75%, 100%) diluted in UltraPure Distilled Water. Tissue was incubated for 60 minutes in each dehydration solution at room temperature. Tissue was stored long-term in 100% Methanol at 4°C. Prior to paraffin processing, tissue was equilibrated in 100% ethanol for an hour followed by 70% ethanol. Tissue was paraffin perfused using an automated tissue processor (Leica ASP300) with 1 hour solution changes overnight. Paraffin processed tissue was embedded and stored at room temperature with silica desiccant packets in a sealed container.

### Multiplex Fluorescent *In Situ* Hybridization (FISH) and protein staining

Paraffin blocks were sectioned to generate 5 µm-thick sections. Sections were used within one week for optimal results, and the assay was carried out in RNase-free conditions by treating all materials with RNase-away (Molecular Bioproducts Inc., 7005-11) prior to use. Slides were stored at room temperature in a sealed slide box with silica desiccant packets. Slides were baked for 1 hour in a 60°C dry oven a day prior to starting the procedure. The multiplex fluorescent *in situ* hybridization (FISH) protocol was performed according to the manufacturer’s instructions (ACD; RNAscope Multiplex Fuorescent v2 manual protocol, 323100-USM) under standard antigen retrieval conditions and optimized protease treatment conditions for each tissue (lung 4 min., intestine 30 min., kidney 20min, HIO 20 min). Immunofluorescent protein staining was performed as previously described (Spence et al., 2009) on 5-7 µm sections. A list of RNAscope probes, TSA reagents, and antibodies can be found in the Key Resources Table. Each FISH stain was performed on at least 2 biological replicates. All imaging was done using a Leica SP5 or Nikon A1 confocal and images were assembled using Photoshop CC. Imaging parameters were kept consistent for all images within the same experiment and any post-imaging manipulations (i.e. brightness, contrast, LUTs) were performed equally on all images from a single experiment.

### Wholemount staining of HIOs

HIOs were fixed for 30-60 minutes depending on age in 10% NBF at room temperature on a rocker. Samples were washed three times for 30-minutes in blocking solution (5% normal donkey serum in PBS with 0.1% triton). HIOs were transferred to permeabilization solution (PBS with 0.25% triton-spheroids or PBS with 0.5% triton-HIOs) for 30-60 minutes at room temperature, followed by three 30-minute washes in blocking solution and an additional 1 hour incubation in blocking solution. Primary antibodies against E-cadherin and CD144 were added to blocking solution and incubated for 24 hours at 4 °C (see key resources table for details). The following day, primary antibodies were removed and HIOs were subjected to three 30-minute washes in blocking solution. Appropriate fluorophore conjugated secondary antibodies were incubated with samples for 24 hours at 4 °C. The next day, secondary antibodies were removed and HIOs were washed three times for 30-minutes in blocking solution. DAPI (0.1 mg/ml) was added to the first. Samples were mounted onto slides containing secure-seal spacers (Invitrogen, S24737). Optical clearing was achieved by incubating HIOs in Focus Clear (CelExplorer, FC-101) for 10-20 minutes at room temperature. This process was repeated with fresh Focus Clear until tissue was cleared. Focus Clear was replaced by Prolong Gold and slides were coverslipped. All imaging was done using a Nikon A1 confocal and images were assembled using Photoshop CC. Z-stack series (∼0.8-1.25 µm steps) were captured and 3D rendering was performed using Imaris.

### Isolation of endothelial cells from primary tissue

Primary human fetal tissue was dissociated into single cell suspensions for FACS isolation of ECs according to a previously published protocol (van Beijnum et al., 2008). Single cell suspensions were passed through a 70µm filter, pelleted, and resuspended in staining buffer comprised of PBS with 1x PenStrep, 2 µM EDTA, and 2% FBS. Cells were stained with CD144-APC (0.88 µg/ml) and CD31-PE (0.88 µg/ml) or corresponding isotype controls (0.88 µg/ml) for 30 minutes on ice in a total volume of 100 µl/ per 1×10^7^ cells. Cells were washed three times in excess staining buffer accompanied by centrifugation at 300xg for 5 mins between washes. Cells were resuspended in staining buffer with DAPI (0.2 µg/ml). Antibody stained samples and controls were analyzed and sorted on a FACSAria III cell sorter (BD), and analysis was performed using BD FACSDiva software. Any post-acquisition analysis was performed using FlowJo. Cells were sorted into staining buffer, and after sorting cells were snap frozen and stored at −80 °C prior to RNA isolation.

### Flow cytometric analysis of HIOs

HIOs were removed from Matrigel droplets and transferred to an enzymatic solution. The enzyme solution is comprised of 1 ml dispase (2.5 units/mg, Gibco, 17105-041) and 9 ml collagenase type II (0.1%,Gibco, 17101-015) in PBS per 1 gram of tissue. Tissue digestions were carried out (∼1 hour) at 37 °C, agitating the solution every 10 minutes via stereological pipetting. Enzymatic reactions were quenched by adding 2x the volume of serum-containing (20%) FBS media (DMEM:F12). Cell suspensions were then passed through a 70 µm filter to remove any undigested clumps. Cells were spun down (400 g for 5 mins at 4 °C) and resuspended in a staining buffer comprised of PBS with 1x PenStrep, and 2% FBS. Cells were counted using a hemacytometer, spun down (300 g for 10 minutes), and resuspended in appropriate volumes (100-200 µl) for antibody staining. Cell suspensions were stained with CD31-APC, CD144-PE, or corresponding isotype controls according to the manufacturers recommended dilution (see key resources table). Staining took place at 4 °C for 10 minutes. Cell suspensions were washed by adding 2 mls of buffer, followed by centrifugation (300 g for 10 minutes). Pellets were resuspended in 500 µl of buffer, and DAPI (0.2 µg/ml) was added to appropriate staining conditions. Flow cytometric analysis was performed using a Sony MA900 cell sorter and accompanying software.

### RNA isolation and Bulk RNAseq of primary ECs

RNA was isolated from snap frozen cell pellets using the RNeasy Mirco Kit (74004, Qiagen), according to manufacturer’s guidelines. cDNA libraries were prepared using the SMARTer Stranded Total RNA-Seq Kit v2-Pico Input (634412, Takara). A total of 32 samples were sequenced for 50-bp single-end reads across 4 lanes on an Illumina HiSeq 2500 by the University of Michigan Advanced Genomics Core. Bulk RNA sequencing analysis was performed as previously descried (Tsai et al., 2018). All reads were aligned to an index of transcripts from human genes within the Ensembl GRCh38 and quantified using Kallisto (Bray et al., 2016). Gene level data generated from Kallisto was used for TMM normalization in edgeR to create normalized data matrix of pseudocounts (Robinson et al., 2010). Principal component analysis and sample clustering were done in R using the ‘cluster’ and Bioconductor ‘qvalue” packages (Storey et al., 2019). Genes were clustered by k-means clustering, using the KMeans function of the scikit learn package, after z-score transformation of pseudocount data (Pedregosa et al., 2011).

### Single Cell Preparation of tissue for single cell RNA sequencing

#### Human Fetal Tissue

Cell dissociations were carried out similar to previously published methods (Miller et al., 2020). To dissociate human fetal tissue to single cells, tissue was mechanically minced into small fragments, and in a petri dish filled with ice-cold 1X HBSS (with Mg^2+^, Ca^2+^). This tissue was then transferred to a 15 mL conical tube. Dissociation enzymes and reagents from the Neural Tissue Dissociation Kit (Miltenyi, cat. no. 130-092-628) were used, and all incubation steps were carried out in a refrigerated centrifuge pre-chilled to 10°C unless otherwise stated. All tubes and pipette tips used to handle cell suspensions were pre-washed with 1% BSA in HBSS to prevent adhesion of cells to the plastic. Tissue was treated for 15 minutes at 10°C with Mix 1. Mix 2 was added to the digestion, and tissue was incubated for 10 minute increments at 10°C until digestion was complete. After each 10 minute incubation, tissue was agitated using a P1000, and tissue digestion was visually assessed under a stereo microscope. This process continued until the tissue was fully digested. Cells were filtered through a 70 µm filter coated with 1% BSA in 1X HBSS, spun down at 500g for 5 minutes at 10°C and resuspended in 500µl 1X HBSS (with Mg^2+^, Ca^2+^). 1 mL Red Blood Cell Lysis buffer (Roche cat. No 11814389001) was then added to the tube and the cell mixture was placed on a rocker for 15 minutes at 4°C. Cells were spun down (500g for 5 minutes at 10°C), and washed twice by suspension in 2 mLs of HBSS + 1% BSA followed by centrifugation. Cells were counted using a hemocytometer, then spun down and resuspended (if necessary) to reach a concentration of 1000 cells/µL and kept on ice. Single cell droplets were immediately prepared on the 10x Chromium according to manufacturer instructions at the University of Michigan The Advanced Genomics Core, with a target of capturing 5,000-10,000 cells. Single cell libraries were prepared using the Chromium Next GEM Single Cell 3’ Library Construction Kit v3.1 according to manufacturer instructions.

#### Organoids

To dissociate human intestinal organoids to single cells, organoids were mechanically isolated from Matrigel droplets and then tissue minced into small fragments using a scalpel in a petri dish filled with ice-cold 1X HBSS (with Mg^2+^, Ca^2^). This tissue was then transferred to a 15 mL conical tube. Dissociation enzymes and reagents from the Neural Tissue Dissociation Kit (Miltenyi, cat. no. 130-092-628) were used, and all incubation steps were carried out in a 37°C incubator unless otherwise stated. All tubes and pipette tips used to handle cell suspensions were pre-washed with 1% BSA in HBSS to prevent adhesion of cells to the plastic. Tissue was treated for 15 minutes at 37°C with Mix 1. Mix 2 was added to the digestion, and tissue was incubated for 10 minute increments at 37°C until digestion was complete. After each 10 minute incubation, tissue was agitated using a P1000, and tissue digestion was visually assessed under a stereo microscope. This process continued until the tissue was fully digested. Cells were filtered through a 70 µm filter coated with 1% BSA in 1X HBSS, spun down at 500g for 5 minutes at 4°C and resuspended in 500µl 1X HBSS (with Mg^2+^, Ca^2+^). Cells were counted using a hemocytometer, then spun down and resuspended (if necessary) to reach a concentration of 1000 cells/µL and kept on ice. Single cell droplets were immediately prepared on the 10x Chromium according to manufacturer instructions at the University of Michigan The Advanced Genomics Core, with a target of capturing 5,000-10,000 cells. Single cell libraries were prepared using the Chromium Next GEM Single Cell 3’ Library Construction Kit v3.1 according to manufacturer instructions.

### Quantification and Statistical Analysis

All graphs and statistical tests were performed in GraphPad Prism 8 software. To determine significance differences across multiple groups, a one-way Analysis of Variance (ANOVA) was performed followed by Tukey’s multiple comparisons analysis comparing the mean of each group to the mean of every other group. A p-value of less than 0.05 was considered significant. On graphs, p-values for multiple comparisons after ANOVAs are reported as followed: **** p<0.0001.

### Data preprocessing Cluster Identification and Cell Type Scoring

All single-cell RNA-sequencing was performed with an Illumina Novaseq 6000 at the University of Michigan Advanced Genomics Core. Raw data was processed using the 10x Genomics Cell Ranger v2.1.1-2.2.1 pipeline using human reference genome (hg19) to generate gene expression matrices. Analysis was performed using the Single Cell Analysis for Python toolbox described in previously (Wolf et al., 2018). To ensure input data were of high quality, filtering parameters for gene count range, unique molecular identifier (UMI) counts, and mitochondrial transcript fraction were imposed on each data set. After organ-specific quality filtering parameters were applied, all primary human data sets were combined for the remainder of preprocessing. Gene expression levels were log normalized, highly variable genes were extracted, and effects of UMI count and mitochondrial transcript fraction variations were regressed out by linear regression. Gene expression values were z-transformed before samples were again separated by organ for downstream analysis. A graph-based clustering approach was performed using the top 10-11 principal components. Further dimensional reduction was done using the UMAP algorithm (McInnes et al., 2018), and cluster identification was performed as previously described (Blondel et al., 2008). In certain analyses, endothelial cell clusters were identified based on expression of canonical markers (i.e. *CDH5, KDR*) and computationally extracted and re-clustered.

For cell type scoring of major cell classes (epithelium, mesenchyme, endothelial, neuronal, immune), gene sets for each class were curated in previously published work (Czerwinski et al., 2020; Miller et al., 2020) (Supplementary Table 2), and log normalized and z-transformed raw counts were summed to generate cell type scores. For visualization of these data, cell type scores were mapped onto UMAP embeddings. For organ-specific EC gene signatures (lung EC signature (lECs); intestinal EC signature (iECs); kidney EC signature (kECs)) gene sets from the bulk RNAseq k-means clustering analysis were used. A given gene set was filtered to include the most highly enriched genes using genes with a log normalized z-score ≥ 1.8. This filtering resulted in gene lists containing 137 (lung), 89 (intestine), and 50 (kidney) genes, respectively (Supplementary Table 2). For comparison of primary fetal EC gene sets to HIO ECs, HIO ECs were queried for expression of genes from each set, and the mean log normalized and z-transformed raw counts for each gene signature set was determined (i.e. the mean expression of the lECs, iECs, and kECs sets were determined in HIO ECs). Cell scores were mapped onto the HIOEC embeddings.

## DATA AND CODE AVAILABILITY

The human fetal lung scRNAseq data used for our analyses have been previously published (Miller et al., 2020). The human fetal intestine (Czerwinski et al., 2020) and kidney data deposition is in progress. **Code is available at:** https://github.com/jason-spence-lab/Holloway2020

---

**Supplemental Figure 1.**
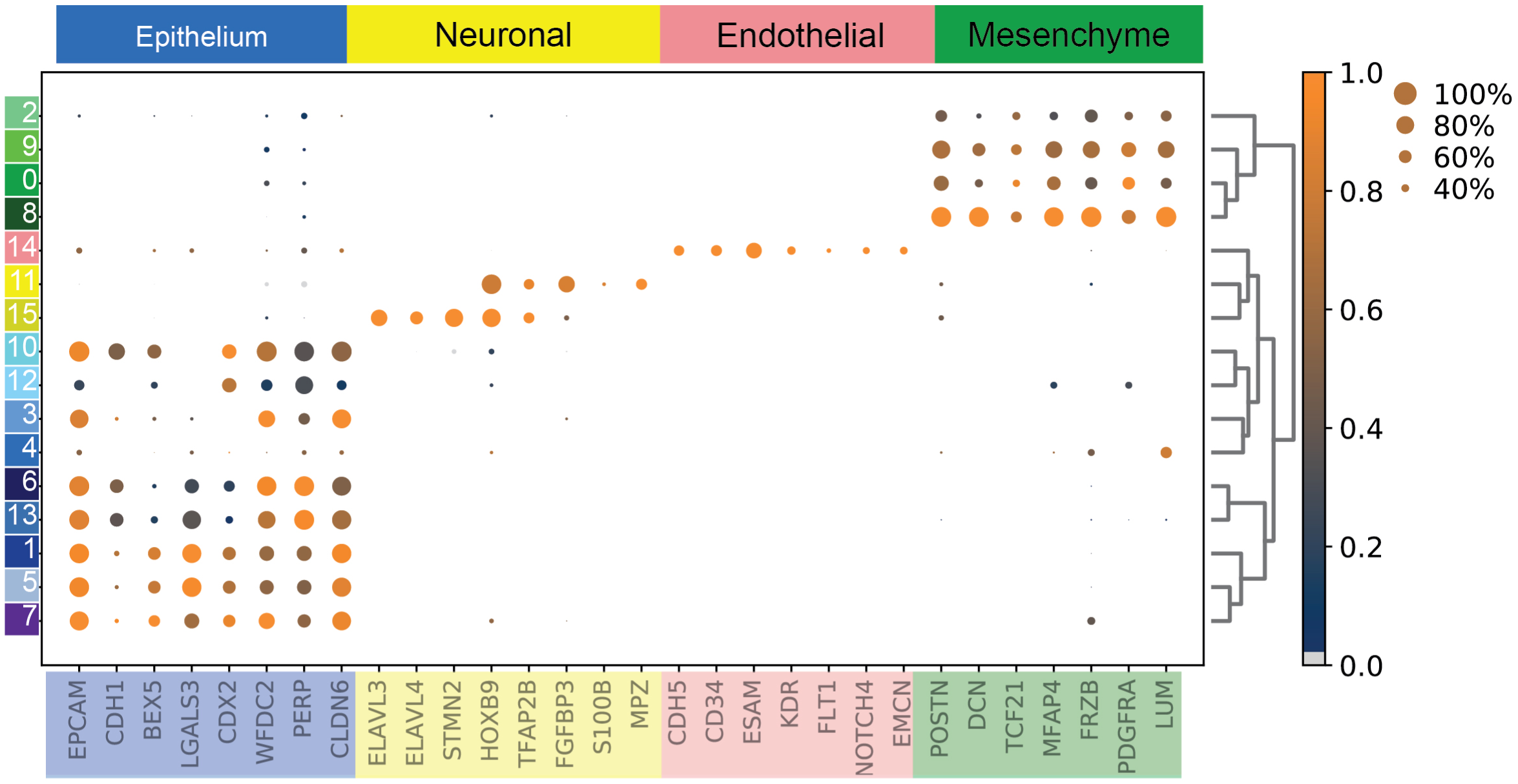
Cell type classification of HIOs. Dotplot of genes associated with major cell classes: epithelial, mesenchymal, endothelial, neuronal genes. Dot size represents the proportion of cells in each cluster expressing a given marker, while color indicates log normalized expression level. Clusters are colored by class assignment: Blue – epithelium; Yellow – neuronal; Red – endothelial; Green – mesenchyme.

**Supplemental Figure 2.**
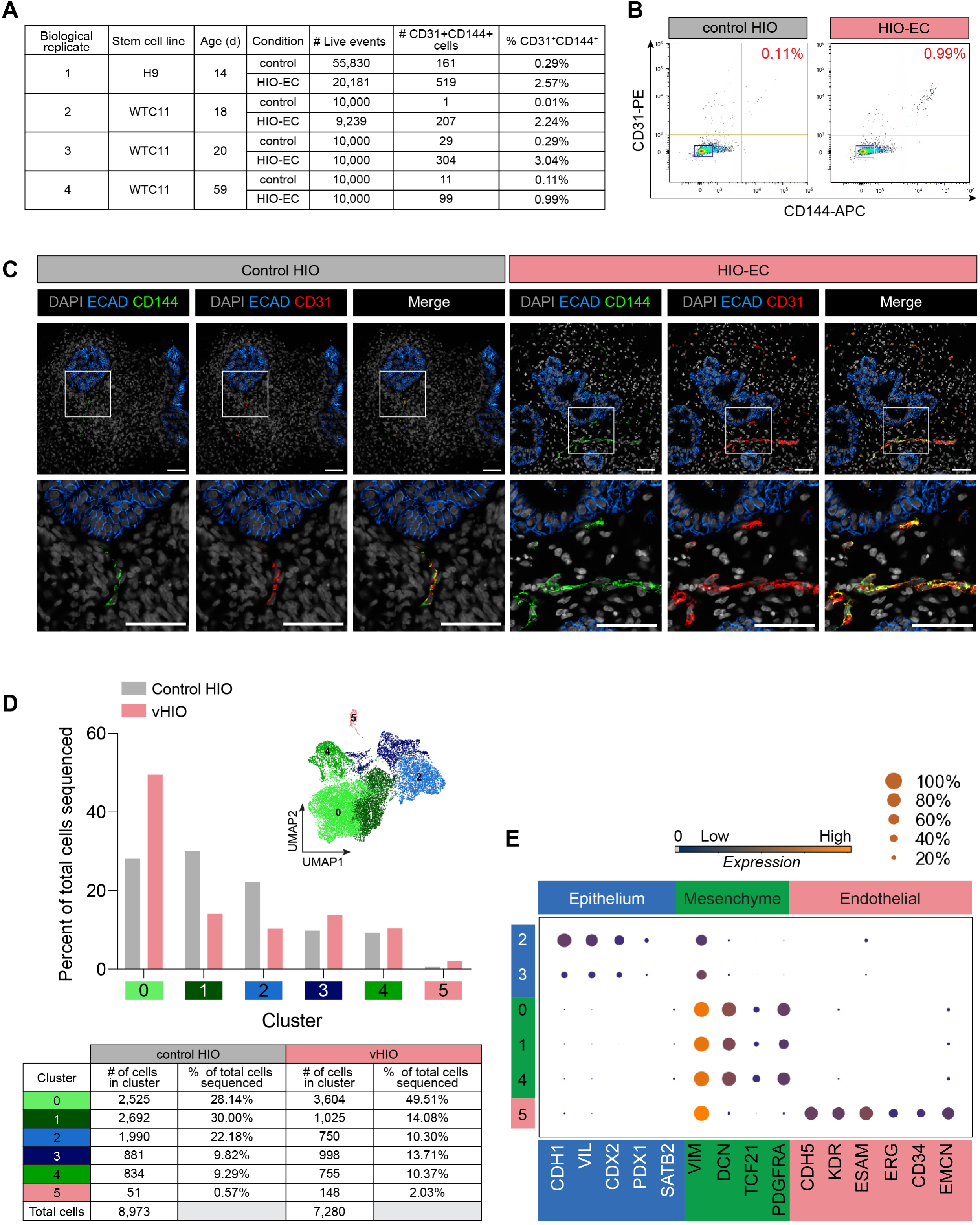
Characterization of vHIOs. **(A)** Summary table for flow cytometric analysis of four independent experiments using two different hPSC lines analyzing CD31+CD144+ and CD31-CD144-cells in matched (i.e. differentiated at the same time) control HIOs and vHIO. **(B)** Flow cytometric analysis from 59d control HIO and vHIO (WTC11 iPSC line) for EC markers CD31 and CD144. **(C)** Maximum intensity projections images from z-series imaging (3-5 µm) of immunofluorescent stains carried out on thin paraffin sections for EC markers CD31(red), CD144 (green), ECAD (blue), DAPI (grey) in batch-matched 22d control and vHIOs. Scalebars represent 50 µm. (D) Single cell RNA-seq analysis of 59d WTC11 iPSC-derived control HIOs and vHIO. Inset UMAP plot shows predicted clusters, and the proportion of cells from control HIOs or vHIOs contributing to each cluster is shown (bar plot, table). (E) Dotplot assigning cell classes to control HIOs and vHIOs: intestinal epithelium (*CDH1, VIL, CDX2*), mesenchymal (*VIM, DCN, TCF21, PDGFRa*), and endothelial (*CDH5, KDR, ESAM, ERG, CD34, and EMCN*) markers. Additionally, HIOs were patterned into proximal (duodenal) small intestine as determined by expression of *PDX1*, and low-to-absent expression of the distal-small intestinal (ileum) and colonic epithelial marker *SATB2*. Dot size represents the proportion of cells in each cluster expressing a given marker, while color indicates log normalized expression level.

**Supplemental Figure 3.**
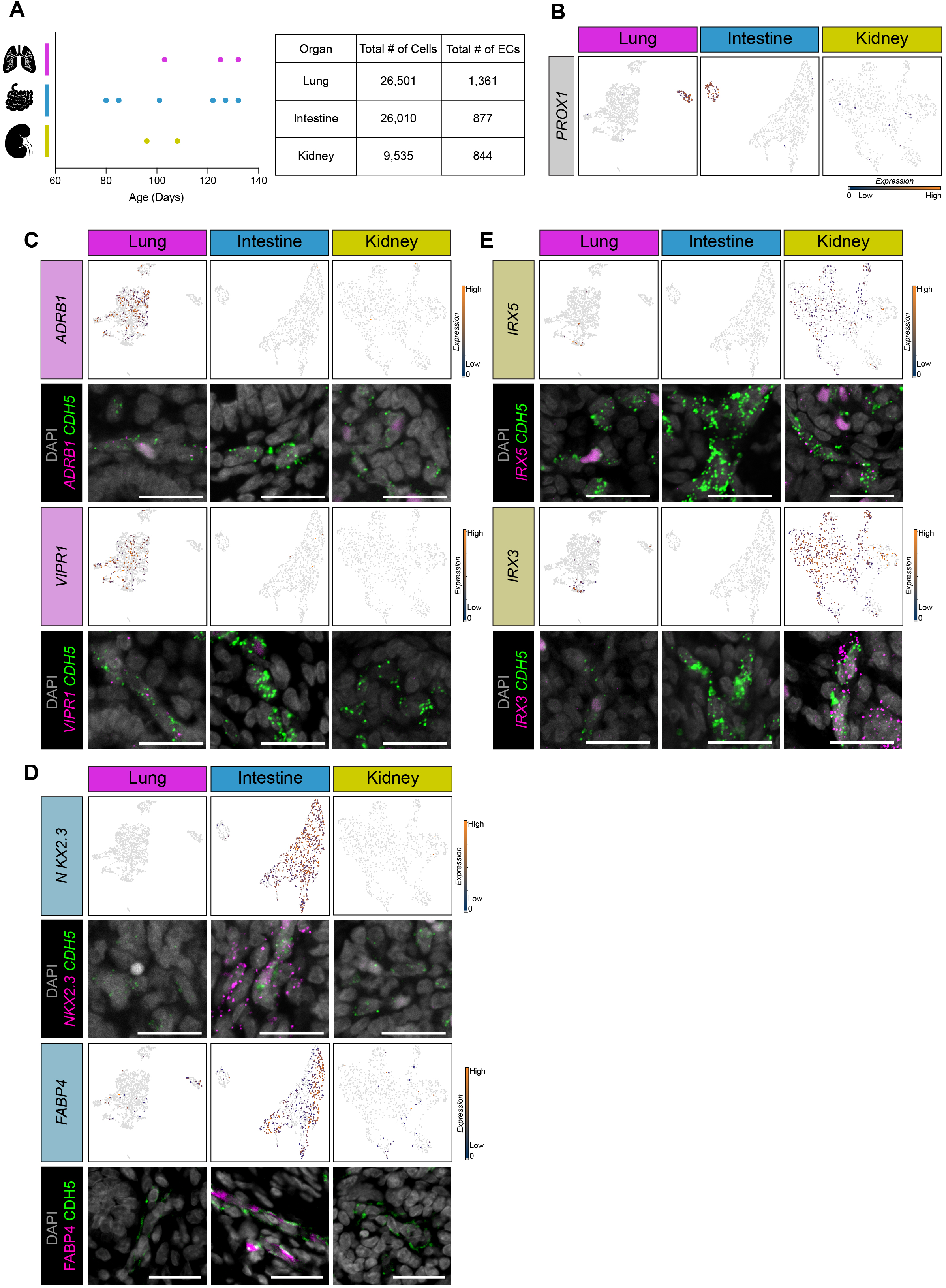
Additional scRNAseq and FISH validation of organ-specific EC-enriched signatures. (A) Summary of samples profiled by scRNAseq, along with the total number of cells and the number of ECs profiled for each organ (table). (B) The *CDH5+* EC cluster from each organ was extracted and reclustered (Figure 4A), and feature plots for the lymphatic marker *PROX1* is shown. (C) Feature plots showing expression of lung-specific candidates *ADRB1* and *VIPR1* among primary lung, intestine or kidney ECs with corresponding multiplexed FISH in all three tissues (101d) with these markers (pink), pan-EC marker *CDH5* (green), and DAPI (grey). (D) Feature plots depicting expression of intestine-specific candidates *NKX2.3* and *FABP4* among primary ECs. Corresponding multiplexed FISH for *NKX2.3* (pink) and pan-EC marker *CDH5* (green), and DAPI (grey) in 117d human. Immunofluorescent staining for FABP4 (pink), CDH5 (pink), and DAPI (grey) in 120d tissue. (E) Feature plots depicting expression of kidney-specific candidates *IRX5* and *IRX3* among primary ECs with corresponding multiplexed FISH with these markers (pink), pan-EC marker *CDH5* (green), and DAPI (grey) in 117d tissue. Note: large pink areas in E correspond to background from red blood cells; specific signal is seen as small bright magenta spots.

**Supplemental Figure 4.**
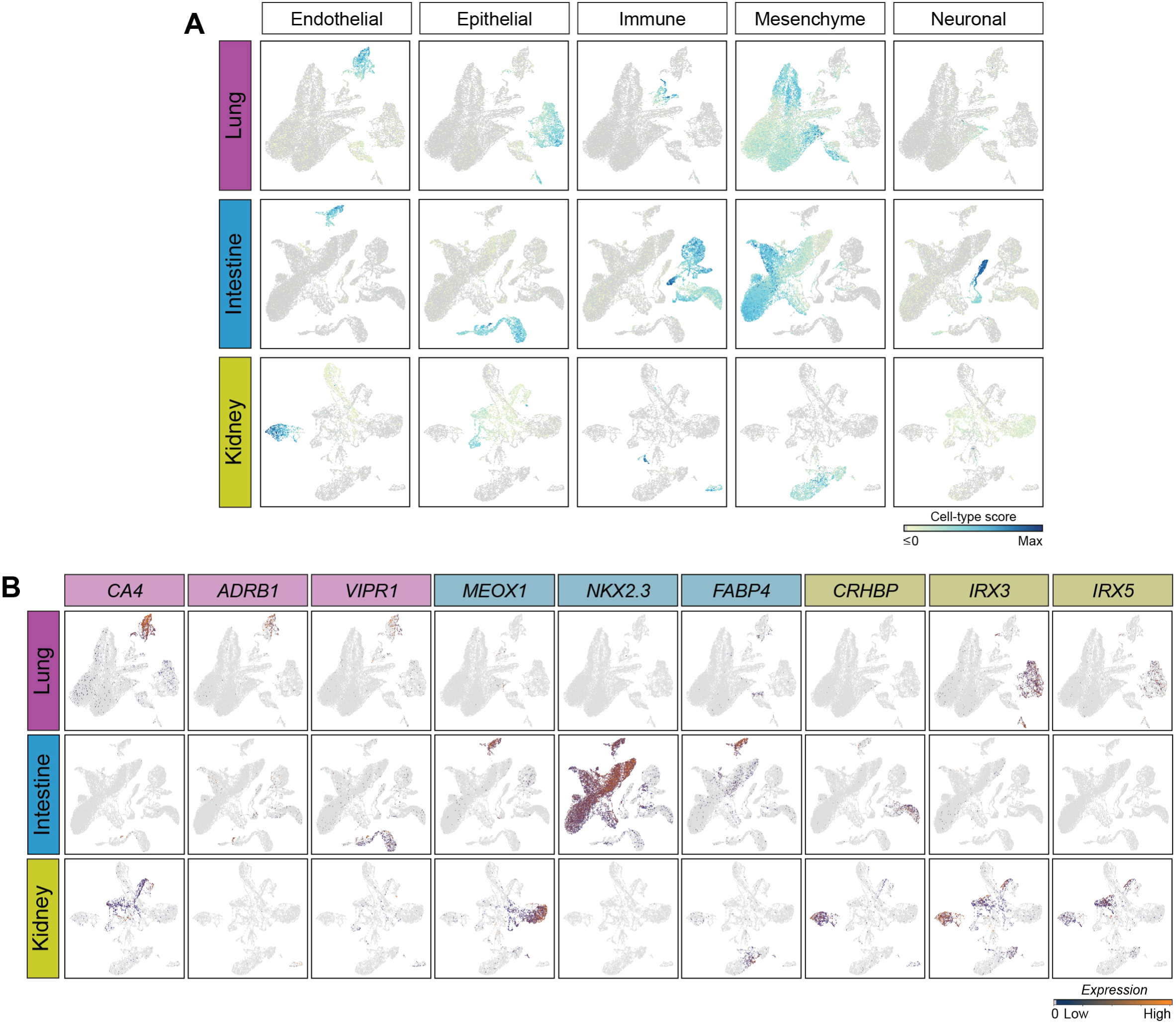
Organ-specific EC gene marker expression across organs and major cell classes. (A) Cell type scoring for major cell classes in fetal lung, intestine and kidney data sets, determined as the summed normalized gene expression of canonical markers for endothelial, epithelial, mesenchymal, immune, and neuronal cell populations. B) Feature plots showing expression of organ-specific EC enriched genes across all cells sequenced for each organ. This analysis shows that a given gene may be specific to the ECs of only one organ (i.e. *IRX3* in kidney ECs), but that this gene may still be expressed in non-ECs of another organ (i.e. *IRX3* is also expressed in lung epithelium).

**Supplemental Figure 5.**
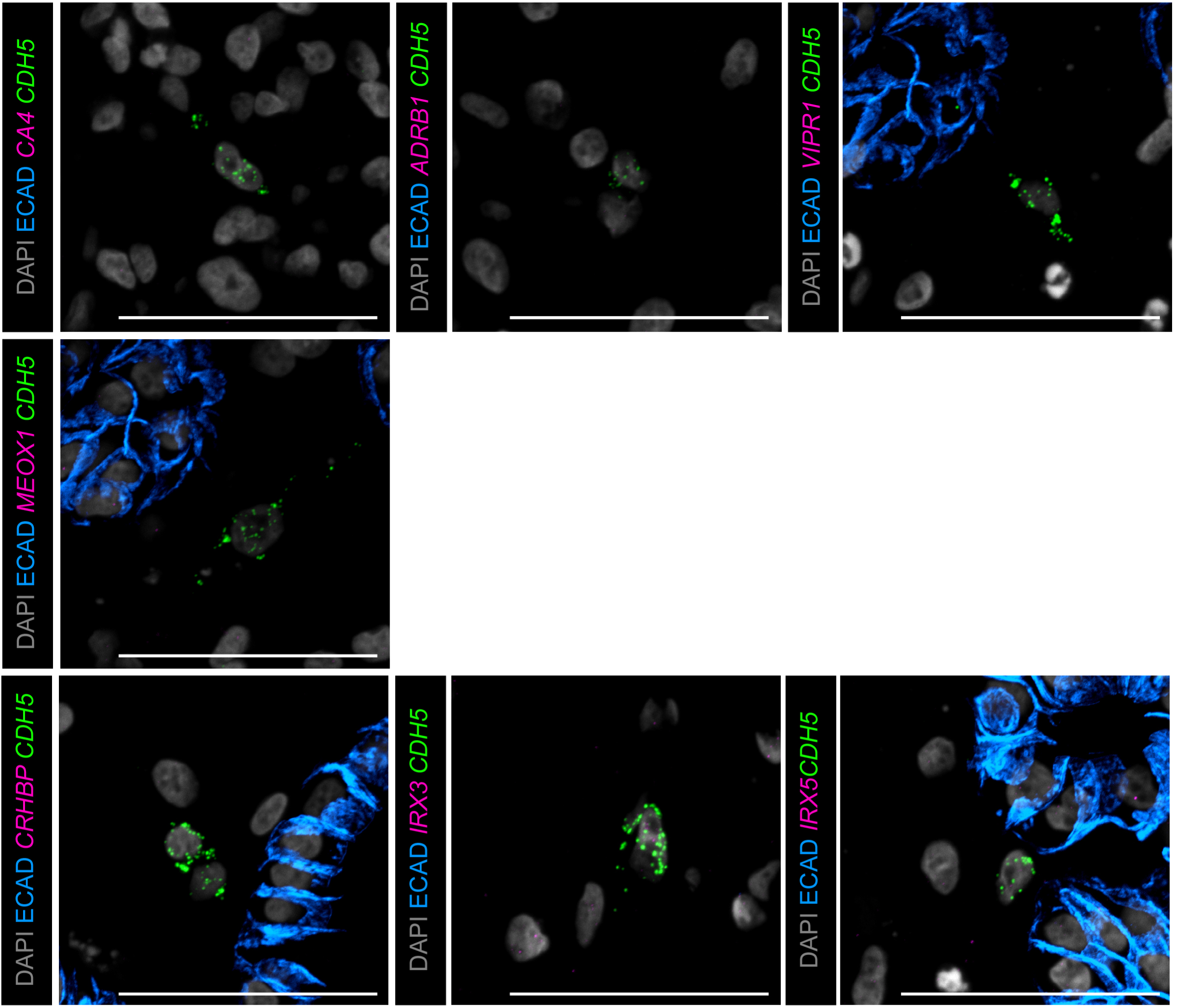
Organ-specific EC gene marker expression in vHIOs after 29d in culture. Top row: FISH of the lung EC markers *CA4, ADRB1, VIPR1* (pink) co-stained with the pan-EC marker *CDH5* (green), protein staining for ECAD (blue), and DAPI (grey) in 29d vHIOs. Middle row: FISH of the intestine EC marker *MEOX1* (pink) co-stained with the pan-EC marker *CDH5* (green), protein staining for ECAD (blue), and DAPI (grey) in 29d vHIOs. Bottom row: FISH of the kidney EC markers *CRHBP, IRX3, IRX5* (pink) co-stained with the pan-EC marker *CDH5* (green), protein staining for ECAD (blue), and DAPI (grey) in 29d vHIOs. These data show that all markers were absent from 29d vHIOs. Scalebars represent 50 µm.

